# Metabolic signatures of cone-dominant and cone-degenerating retinas

**DOI:** 10.64898/2026.08.11.744269

**Authors:** David S. Hansman, Tuan Ngo, Jeff Hao, Isabella Mascari, Mark Eminhizer, Hongwei Ma, Saira Rizwan, Audrey Anderson, Artjola Puja, Jinyu Lu, Qingyan Wang, Ying Zhang, Yinxiao Xiang, Cloe Ratliff, Diana Alabdallat, Xi-Qin Ding, Jianhai Du

## Abstract

Cone photoreceptors are essential for daylight vision, and their degeneration has more profound visual consequences than rod loss in retinal degenerative diseases. Metabolic dysfunction is closely associated with cone degeneration, however the relatively small population of cones in mice and humans have limited our understanding of cone-specific metabolism.

Here, we leveraged cone-dominant and cone-degeneration mouse models including *Nrl^-/-^*, *Cnga3^-/-^*, and high-dose Triiodothyronine (T3) treatment to investigate cone-specific metabolism and their metabolic impacts on the retinal pigment epithelium (RPE). Across models, cone-dominant retinas consistently showed lower pyruvate abundance alongside increases in glutathione, purines, pentose phosphate pathway intermediates, and one-carbon metabolites. Increases in several key amino acids were also associated with higher cone abundance, such as proline, arginine, alanine, valine, leucine, and hypotaurine. Strikingly, aminoadipate, an intermediate in lysine catabolism, was the most robustly changed metabolite in the retina, showing highly consistent increases across models.

Relative cone increases were also associated with metabolic changes in the RPE/choroid. Like the retina, RPE/choroids showed consistent increases in aminoadipate, proline, and hypotaurine, as well as xanthosine and betaine, alongside decreases in uracil. Moreover, proteomic analysis of *Nrl^-/-^* mice showed decreases in many key metabolite transporters in the RPE/choroid, including carriers for glucose, lactate, aspartate, glutamate, serine, lysine, taurine, and proline. Collectively, these findings further our understanding of cone-specific metabolism and highlight potential cone-specific metabolic vulnerabilities in retinal degeneration.

## 1. Introduction

The retina is the most energy-demanding tissue in the body^1^, largely due to the profound bioenergetic requirements of rod and cone photoreceptors^2,3^. Cones have approximately double the energy demand of rods^4^, requiring more ATP to support Ca^2+^ extrusion via ATP-dependent pumps^4,5^. Reflecting this demand, cones have double the number of mitochondria and threefold more cristae surface area than rods in mice^6^, and several-fold greater mitochondrial volume in non-human primates^7,8^. Unlike rods, cones are resistant to deficiency in aerobic glycolysis enzymes^9^, glucose depravation^10,11^, and glycolytic inhibition^12^. Beyond glucose metabolism, cones also show specialized amino acid and nucleotide uptake and release^13^, membrane lipid composition^14^, and metabolic gene expression^15^. Despite advances in defining cone-specific function and gene expression^15,16^, the relatively small population of cones, particularly in mice and humans^17,18^, has posed a major barrier to characterizing the cone metabolome.

The outer retina exchanges sugars, amino acids, ketone bodies, and other metabolites with the overlying retinal pigment epithelium (RPE) monolayer^13,19–25^. RPE metabolism is essential for retinal homeostasis; RPE-specific metabolic dysfunction induced by activation of hypoxia-inducible signaling^26^, pro-glycolytic mammalian target of rapamycin (mTOR) signaling^27^, mitochondrial impairment^28^, or defective lipid efflux^29^ drives photoreceptor degeneration in genetic mouse models. Conversely, photoreceptor-specific activation of mTOR in mice causes lipid dysregulation, polynucleation, and atrophy in RPE cells^30^. However, cone-specific metabolic interactions with the RPE remain poorly understood.

Dysregulated metabolism is associated with many retinal diseases. Retinitis pigmentosa (RP) models consistently show metabolic gene dysregulation^31^, while constitutive activation of the insulin/mTOR pathway promotes cone survival in RP mice^32^. Moreover, mutations in lipid and mitochondrial metabolism are associated with cone-rod dystrophies - inherited retinal diseases marked by primary cone loss^33–36^. Age-related macular degeneration (AMD) is associated with metabolic dysregulation in the retina and RPE^21,37,38^. Because cone photoreceptors are most densely concentrated in the macula^39,40^, the primary site of AMD pathology, defining cone-specific metabolism may improve our understanding of disease progression. However, cone-specific metabolism remains unclear, and its characterization may reveal therapeutic targets to treat retinal pathologies.

Animal models with altered photoreceptor populations provide valuable tools for investigating cone-specific metabolic features. Neural retina leucine zipper (Nrl) is a transcription factor essential for rod differentiation during retinal development^41,42^. *Nrl^-/-^* mice lack rods entirely and instead develop retinas composed of functional cone-like photoreceptors, making them a widely used model for studying cone-specific biology^41,43,44^. Cyclic nucleotide-gated channel alpha-3 (CNGA3) is a cone-specific ion channel required for phototransduction, mediating Na^+^ and Ca^2+^ influx in response to high intracellular cyclic guanosine monophosphate (cGMP)^45^. Mutations in *CNGA3* are associated with cone dystrophies and loss of color vision in humans^46,47^. *Cnga3^-/-^* mice show complete loss of cone function and early-onset cone degeneration^48,49^. The thyroid hormone triiodothyronine (T3) is a critical regulator of cone maintenance, differentiation, and subtype specification, including in human retinal organoids^50,51^. However, systemic administration of high-dose T3 in mice induces degeneration of both rods and cones, impairing phototransduction and mitochondrial metabolism^52,53^. Especially in combination, these models offer valuable systems to interrogate metabolic asymmetries between rods and cones, as well as their differential interactions with the RPE.

Using these models, we rigorously investigate cone-specific retinal metabolism, and cone-RPE interactions. Strikingly, we find that cone-dominant retinas show consistent increases in amino acids such as aminoadipate, valine, proline, and hypotaurine, alongside large decreases in pyruvate. Proteomic analysis of *Nrl^-/-^* mice reveals that the RPE/choroid responds to increased cones by downregulating key metabolite transporters, including carriers for glucose, lactate, proline, and taurine. These findings refine our understanding of cone metabolism and identify candidate pathways that may be leveraged to preserve cone function in disease.

## 2. Results

### 2.1. Altered amino acid and nucleotide metabolites in cone-dominant retinas

To investigate cone-specific metabolism, we performed targeted metabolomics of 187 metabolites across major pathways in retinas and RPE/choroids from models with varying proportions of cones (Figure 1) (Table S1). Using this data, we performed four comparisons: (1) *Nrl^⁻/⁻^* mice, which lack rods and are instead populated by cone-like cells, compared to *Nrl^⁺/⁻^* controls with normal photoreceptor populations; (2) Wild-type (WT, littermate control) mice compared to *Cnga3^⁻/⁻^* mice, which show early cone dysfunction and degeneration; (3) *Nrl^⁻/⁻^* mice compared to *Nrl^⁻/⁻^ Cnga3^⁻/⁻^* double knockout mice; and (4) untreated *Nrl^⁻/⁻^* mice compared to *Nrl^⁻/⁻^*mice treated with T3 (Table S2-S8).

**Figure 1.**
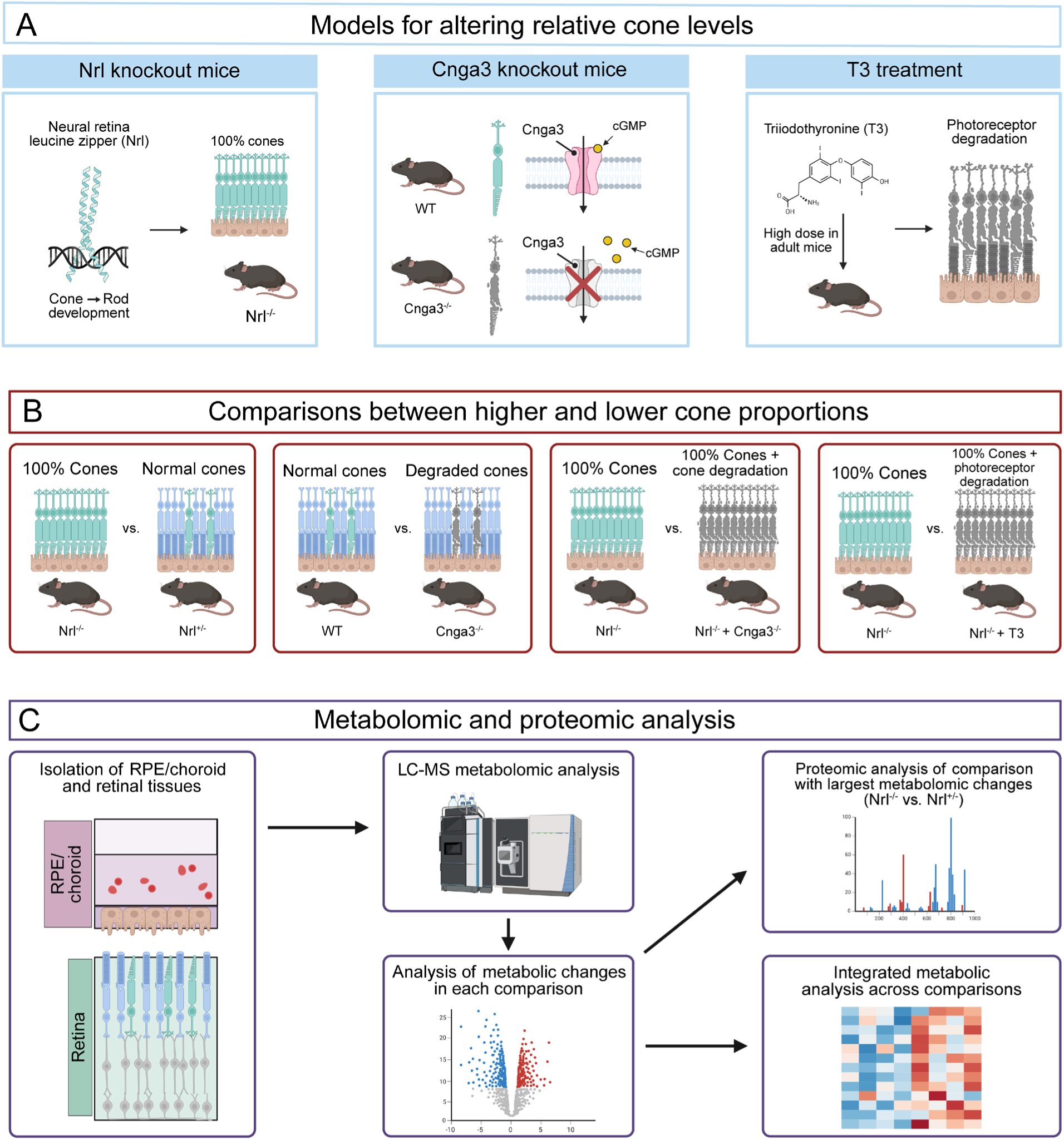
Models of altered cone levels and experimental methodology. Nrl is a transcription factor that promotes the differentiation of retinal precursor cells into rod photoreceptors (A). *Nrl^-/-^* mice lack rods and instead develop outer retinas composed of cone-like photoreceptors. Cnga3 is a cone-specific ion channel involved in phototransduction, and *Cnga3^-/-^* mice show early-onset cone degeneration. Systemic administration of high doses of the thyroid hormone T3 induces degeneration in both rod and cone photoreceptors. The present study uses combinations of these models to perform LCMS-based metabolomic comparisons of retinal and RPE/choroid tissues from mice with relatively high and low cone levels (B). The comparison between *Nrl^-/-^* and *Nrl^+/-^* mice showed the most extensive metabolic changes, so these mice were used for further proteomic analysis of the retina and RPE/choroid (C). Metabolomic changes were integrated across comparisons to identify metabolic features in the retina and RPE/choroid consistently associated with higher cone abundance.

In *Nrl^⁻/⁻^* retinas, more than 30 metabolites were significantly increased compared to *Nrl^⁺/⁻^* controls, while only 4 were decreased (P < 0.05) (Figure 2A-C). The most decreased metabolite was cGMP, while increased metabolites included amino acids (aminoadipate and threonine), purines (guanine and xanthine), and myo-inositol. Comparison of WT vs. *Cnga3^⁻/⁻^* retinas (Figure 2D-F) yielded 10 significantly altered metabolites, including elevated aminoadipate, cytidine, and uridine among the most elevated. In contrast, nucleotides (ATP, cAMP, cGMP), glucose, and the tricarboxylic acid (TCA) intermediate α-ketoglutarate were reduced. *Nrl^⁻/⁻^* vs. *Nrl^⁻/⁻^ Cnga3^⁻/⁻^* retinas showed only a single significantly increased metabolite, aminoadipate (Figure 2G-I) while levels of cGMP, fructose, riboflavin, and 1-methyladenosine were decreased. Compared with T3 treatment (Figure 2J-L), control *Nrl^⁻/⁻^* retinas contained higher levels of multiple metabolites, including amino acids (aminoadipate, proline, arginine, valine), nucleotides (guanine, AMP), and sphinganine.

**Figure 2.**
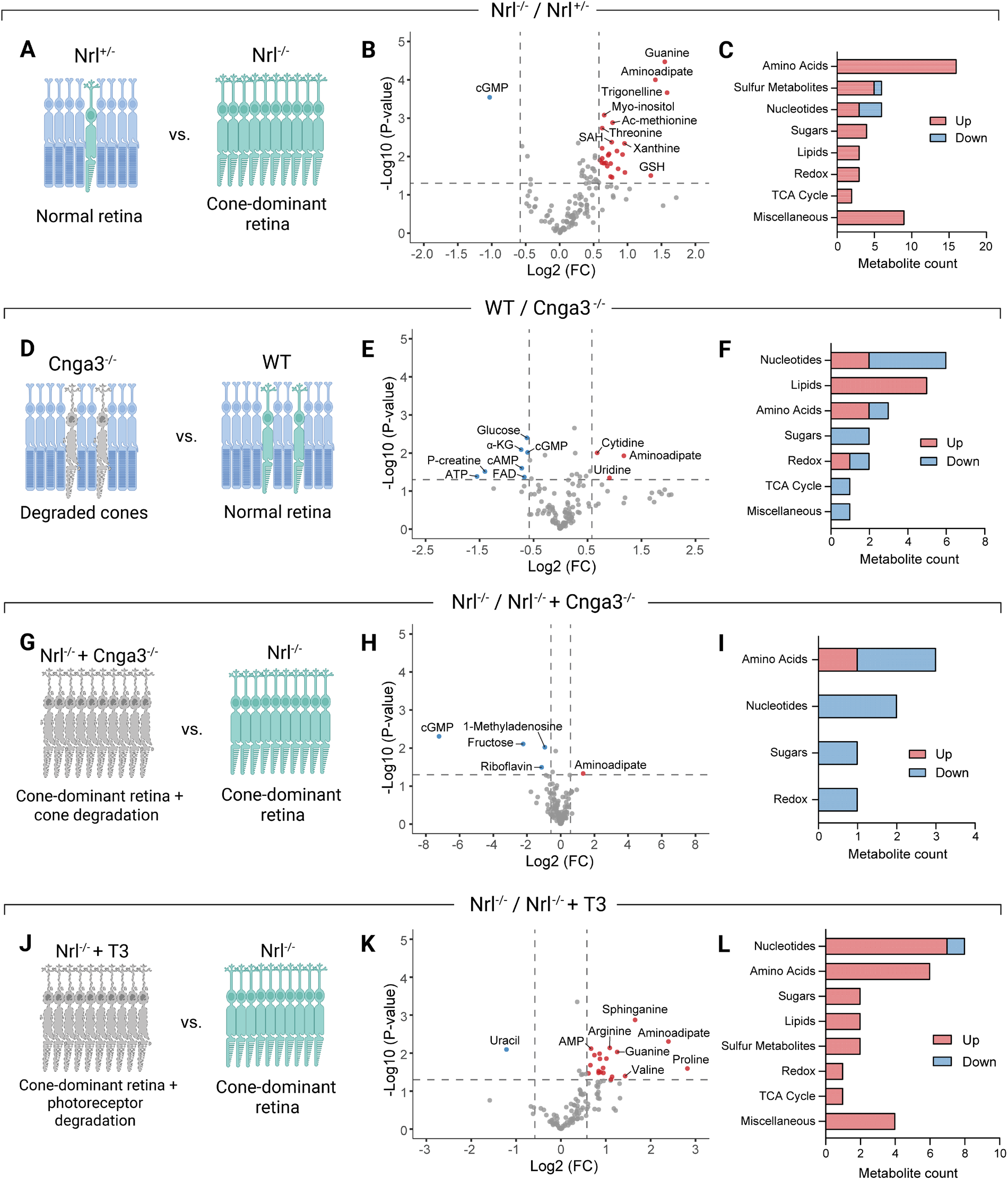
Retinal metabolic changes in models with altered cone levels. Retinal metabolic changes in *Nrl^⁻/⁻^* mice compared to *Nrl^⁺/⁻^* controls (A-C), wild-type compared to *Cnga3^⁻/⁻^* mice (D-F), *Nrl^⁻/⁻^* mice compared to *Nrl^⁻/⁻^ Cnga3^⁻/⁻^* double knockouts (G-I), and *Nrl^⁻/⁻^* mice compared to *Nrl^⁻/⁻^* mice treated with the photoreceptor-degenerating agent T3 (J-L). Volcano plots (B, E, H, K) show retinal metabolite changes in relatively cone-dominant versus cone-deficient mice. Significance thresholds were set at 1.301, corresponding to -log_10_(P < 0.05) as determined using a Student’s unpaired two-tailed T-test. Fold change thresholds were set as > |0.585|, equivalent to metabolites changing by ≥1.5-fold in either direction. Bar charts (C, F, I, L) show classification of significantly changed metabolites in each comparison. Significantly changed (P < 0.05 as determined using a Student’s unpaired two-tailed T-test) metabolites, regardless of the magnitude of fold change, were manually assigned to metabolite categories. Significantly increased and decreased metabolites are colored red and blue, respectively.

### 2.2. Higher cone levels are associated with elevated RPE/choroid amino acids and carnitine metabolites

The RPE engages in complex metabolic interactions with photoreceptors^13,24,54^. To investigate how cones influence RPE metabolism, we also performed targeted metabolomic analysis of RPE/choroid tissue from the relatively cone-dominant and cone-deficient mouse models. RPE/choroid tissues from *Nrl^-/-^* mice contained 17 significantly elevated metabolites compared to control *Nrl^+/-^* mice, including amino acids (hypotaurine, aspartate, and glutamate), carnitine derivatives, and N-methyl-L-proline, while thiamine was the only decreased metabolite (Figure 3A-C). Compared to RPE/choroids in *Cnga3^⁻/⁻^* mice, acetylcarnitine and aminoadipate were the highest among the 6 significantly increased metabolites in WT mice (Figure 3D-F). Conversely, 13 metabolites including cAMP, L-carnosine, citraconic acid, and argininosuccinate were significantly decreased. In the RPE/choroid of *Nrl^⁻/⁻^ Cnga3^⁻/⁻^* double knockout mice, 6 metabolites were significantly decreased compared to *Nrl^⁻/⁻^* controls (Figure 3G-I), including heptadecanoic acid and nucleotides (cGMP, guanine, and inosine). Conversely, proline and kynurenine were the only two significantly increased metabolites, both with only minor increases.

**Figure 3.**
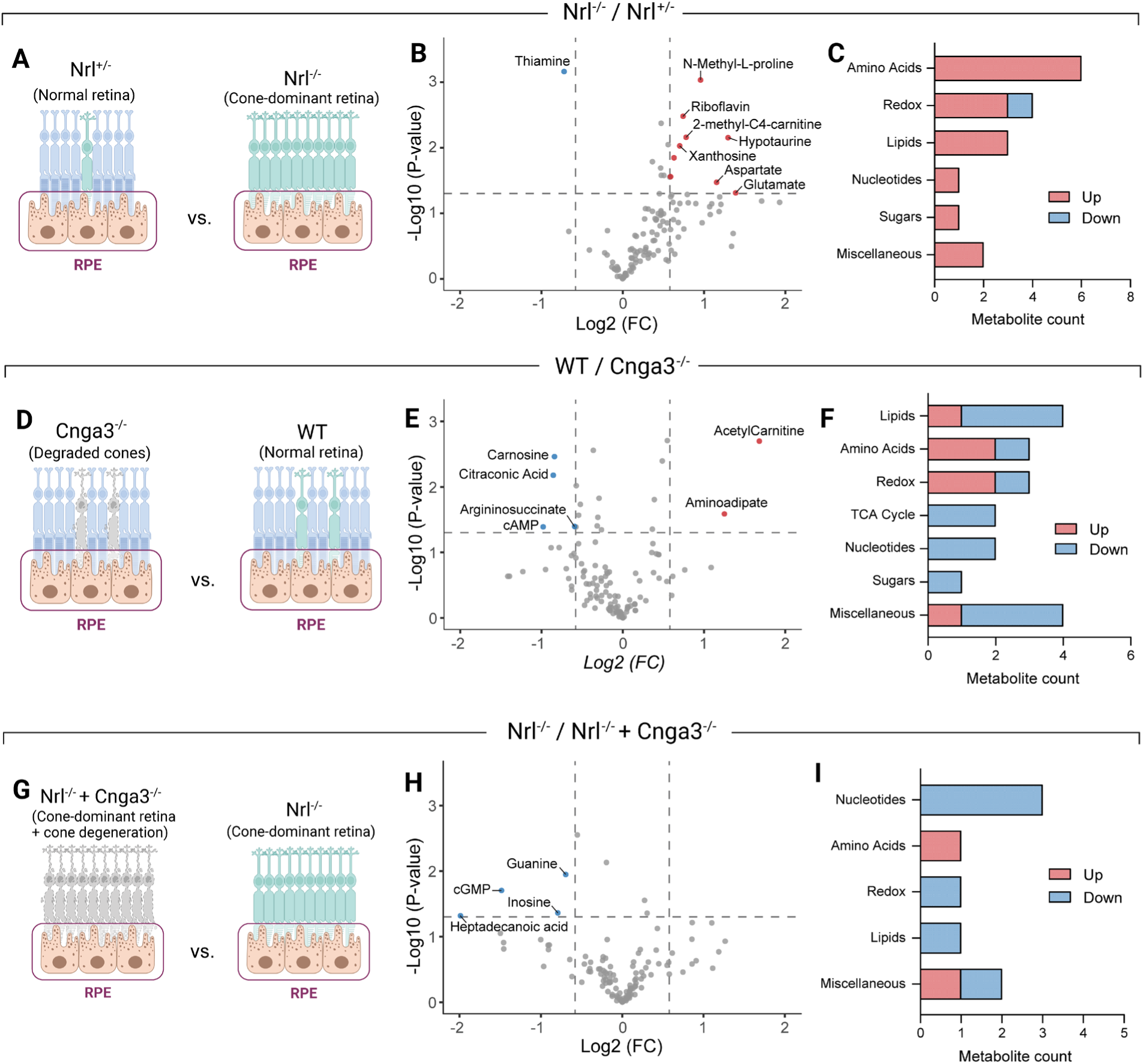
RPE/choroid metabolic changes in models with altered cone levels. RPE/choroid metabolic changes in *Nrl^⁻/⁻^* mice compared to *Nrl^⁺/⁻^* controls (A-C), wild-type compared to *Cnga3^⁻/⁻^* mice (D-F), and *Nrl^⁻/⁻^* mice compared to *Nrl^⁻/⁻^ Cnga3^⁻/⁻^* double knockouts (G-I). Volcano plots (B, E, H) show RPE/choroid metabolite changes in relatively cone-dominant versus cone-deficient mice. Significance thresholds were set at 1.301, corresponding to -log_10_(P < 0.05) as determined using a Student’s unpaired two-tailed T-test. Fold change thresholds were set as > |0.585|, equivalent to metabolites changing by ≥1.5-fold in either direction. Bar charts (C, F, I) show classification of significantly changed metabolites in each comparison. Significantly changed (P < 0.05 as determined using a Student’s unpaired two-tailed T-test) metabolites, regardless of the magnitude of fold change, were manually assigned to metabolite categories. Significantly increased and decreased metabolites are colored red and blue, respectively.

### 2.3. *Nrl^⁻/⁻^* mice show coordinated proteomic and metabolic rewiring across retina and RPE

Among all models, *Nrl^⁻/⁻^*mice showed the most extensive metabolomic changes over *Nrl^+/-^*(Figure S1). Therefore, we performed quantitative proteomics both retina and RPE/choroid to identify altered metabolic proteins in these relatively cone-rich mice (Table S9 and S10).

Differentially expressed protein analysis revealed significant changes in 760 retinal proteins and 140 proteins in the RPE/choroid (Figure 4A-B, Table S9 and S10). As expected, *Nrl^⁻/⁻^* mouse retinas showed lower expression rod-specific proteins, including rhodopsin (RHO), phosphodiesterases PDE6A and PDE6B, transducin alpha (GNAT1), and CNGA1 (Figure 4A, Figure S2). In contrast, several metabolic proteins were upregulated, including myophosphorylase (PYGM), purine nucleoside phosphorylase (PNP), methylenetetrahydrofolate dehydrogenase 1 (MTHFD1), and phosphotriesterase-related protein (PTER) (Figure 4A, Table S11). In the RPE/choroid, higher cone levels were associated with increased cone-specific proteins such as transducin alpha 2 (GNAT2), transducin gamma 2 (GNGT2), opsin 1 short-wave sensitive (OPN1SW), and phosphodiesterase 6C (PDE6C) (Figure 4B, Figure S3), likely derived from phagocytosed outer segments. Notably, many key metabolite transporters were decreased in the RPE/choroids of *Nrl^-/-^* mice, including sodium- and chloride-dependent GABA transporter 2 (GAT2), monocarboxylate transporter 1 (MCT1), solute carrier family 2 member 1 (GLUT1), cationic amino acid transporter 1 (CAT-1), excitatory amino acid transporter 3 (EAAT3), and sodium/imino-acid transporter 1 (SIT1) (Figure 4B, Table S12)

**Figure 4.**
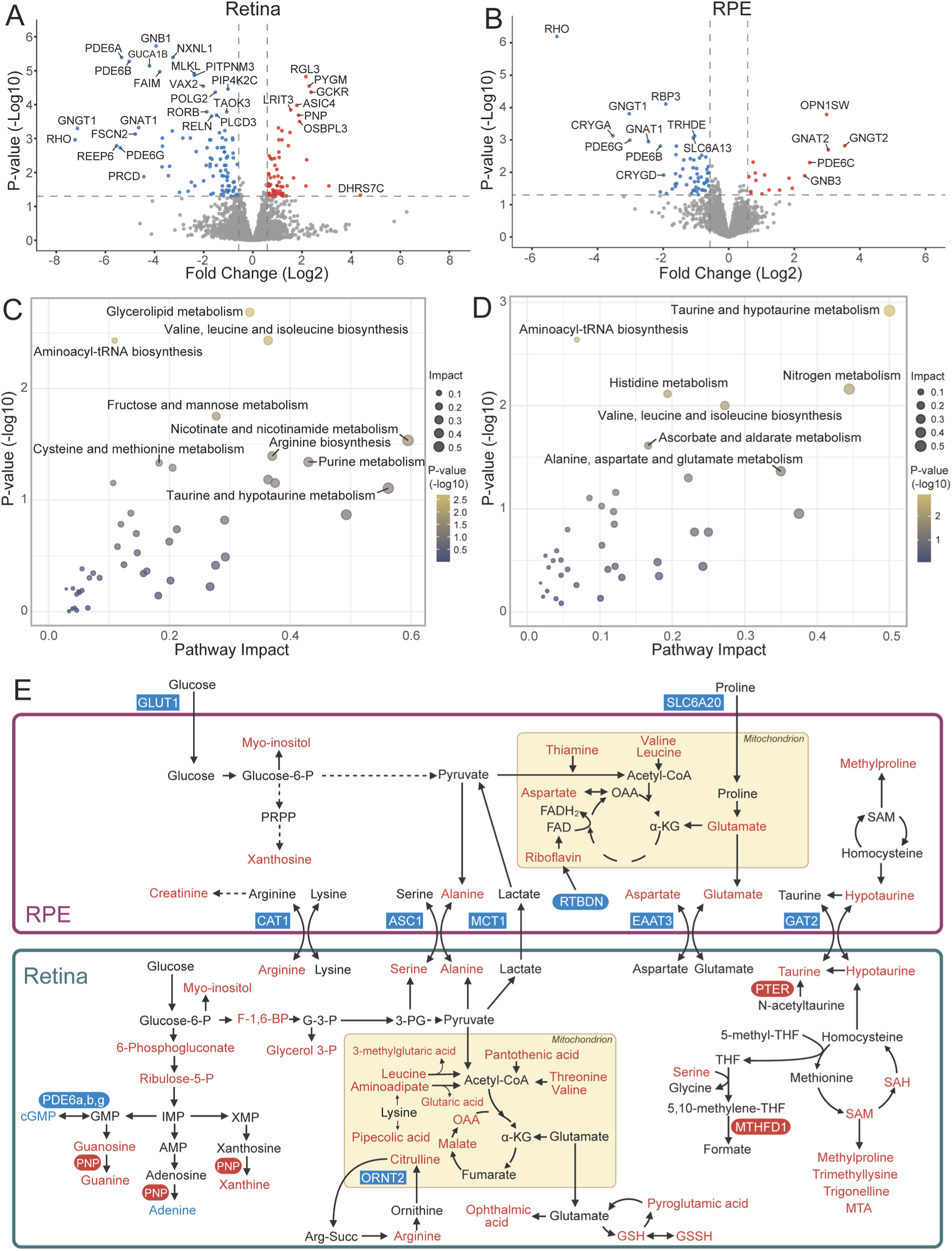
Integrated proteomic and metabolomic analysis of the retina and RPE/choroid in *Nrl^-/-^* versus *Nrl^+/-^* mice. Protein expression in the (A) retina and (B) RPE/choroids in *Nrl^-/-^* relative to *Nrl^+/-^* mice. Significance thresholds were set at 1.301, corresponding to -log_10_(P < 0.05) as determined using a Student’s unpaired two-tailed T-test. Fold change thresholds were set as > |0.585|, equivalent to metabolites changing by ≥1.5-fold in either direction. Integrated pathway analysis of statistically significant (P < 0.05 as determined using a Student’s unpaired two-tailed T-test) protein and metabolite changes in the (C) retina and (D) RPE/choroid between *Nrl^-/-^*and *Nrl^+/-^* mice. Analysis was performed using the joint pathway analysis module on MetaboAnalyst 6.0. (E) Summary diagram of metabolite and metabolic protein changes in the retina and RPE/choroid in *Nrl^-/-^* compared to *Nrl^+/-^* mice.

Integrated pathway analysis of significantly altered metabolites and metabolism-related proteins between *Nrl^⁺/⁻^* and *Nrl^⁻/⁻^* tissues revealed the most affected pathways in the retina included nicotinamide metabolism, purine metabolism, glycerolipid metabolism, and amino acid pathways (taurine/hypotaurine, arginine, leucine, and isoleucine) (Figure 4C). Similarly, in RPE/choroids, relative cone abundance affected nitrogen metabolism and amino acid pathways involving taurine/hypotaurine, alanine, aspartate/glutamate, and branched-chain amino acids (BCAAs) (Figure 4D).

Overall, relative increases in cones were associated with increased abundance of retinal metabolites linked to amino acid catabolism, glutathione regeneration, one-carbon metabolism, and nucleotide degradation (Figure 4E). In contrast, the RPE/choroid showed decreased levels of several important metabolite transporters, including carriers for glucose (GLUT1), lactate (MCT1), proline (SIT1), arginine/lysine (CAT-1), aspartate/glutamate (EAAT3), and taurine/hypotaurine (GAT2).

### 2.4. Integrated meta-analysis shows altered one-carbon, purine, and amino acid metabolism in cone-dominant retinas

To identify metabolites consistently associated with higher cone levels, metabolites were ranked by a ‘consistency score’, a weighted metric incorporating both support (number of comparisons in which a metabolite was detected) and concordance (the consistency of fold-change direction and magnitude across comparisons; see Methods) (Figure 5, Table S13). Among all metabolites, aminoadipate and guanine were identified as the most consistently elevated in cone-dominant retinas. Conversely, cGMP and pyruvate were decreased in 3 out of 4 comparisons, while AMP, proline, and guanosine were consistently increased in the 3 comparisons in which they were detected.

**Figure 5.**
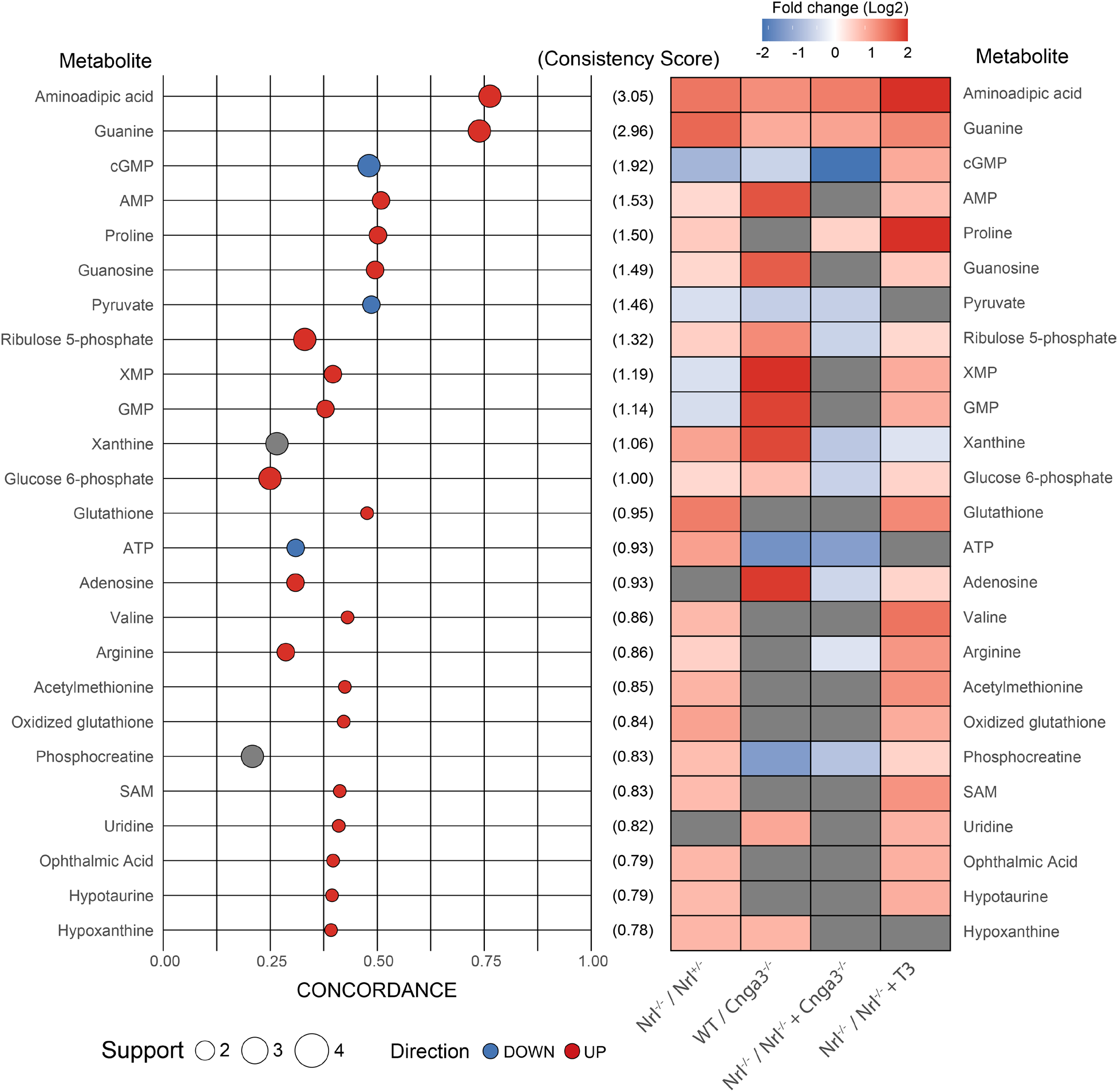
Consistency plot of retinal metabolite changes associated with relative cone abundance. Point size represents support, the number of comparisons in which the metabolite was significantly changed (P < 0.05 as determined using a Student’s unpaired two-tailed T-test). Points are positioned according to their concordance value, a weighted function capturing the consistency and magnitude of change (see methods). Metabolites are ranked based on their consistency score, which is the product of support and concordance values across comparisons. The heatmap shows the log_2_ fold change of metabolites in each comparison. Metabolite changes which were not statistically significant (P > 0.05 as determined using a Student’s unpaired two-tailed T-test) were excluded from the heatmap, as indicated by grey tiles.

To quantify these trends, we calculated Z-scores for metabolites in each comparison and integrated them using Stouffer’s method, generating a meta Z-score for each metabolite (Table S14). Aminoadipate and guanine again showed strong, consistent increases, along with other metabolites such as proline, valine, the one-carbon cycle intermediates SAM and SAH, and several carnitine derivatives (Figure 6A). cGMP and pyruvate were the only metabolites with statistically significant decreases across models. Pathway analysis of significantly altered metabolites (|Z| ≥ 2; *P* < 0.05) revealed enrichment in several key metabolic pathways, including amino acid metabolism (cysteine and methionine, arginine and proline), central carbon metabolism (pyruvate metabolism, TCA cycle), and nucleotide-associated pathways (purine metabolism, PPP) (Figure 6B). These metabolites predominantly fell into categories such as amino acids, nucleotides/nucleosides, and taurine/sulfur-related metabolites (Figure 6C). Overall, cone-dominant retinas were associated with high aminoadipate, proline, myo-inositol, hypotaurine, as well as one-carbon and PPP metabolites, while levels of cGMP and pyruvate were reduced (Figure 6D).

**Figure 6.**
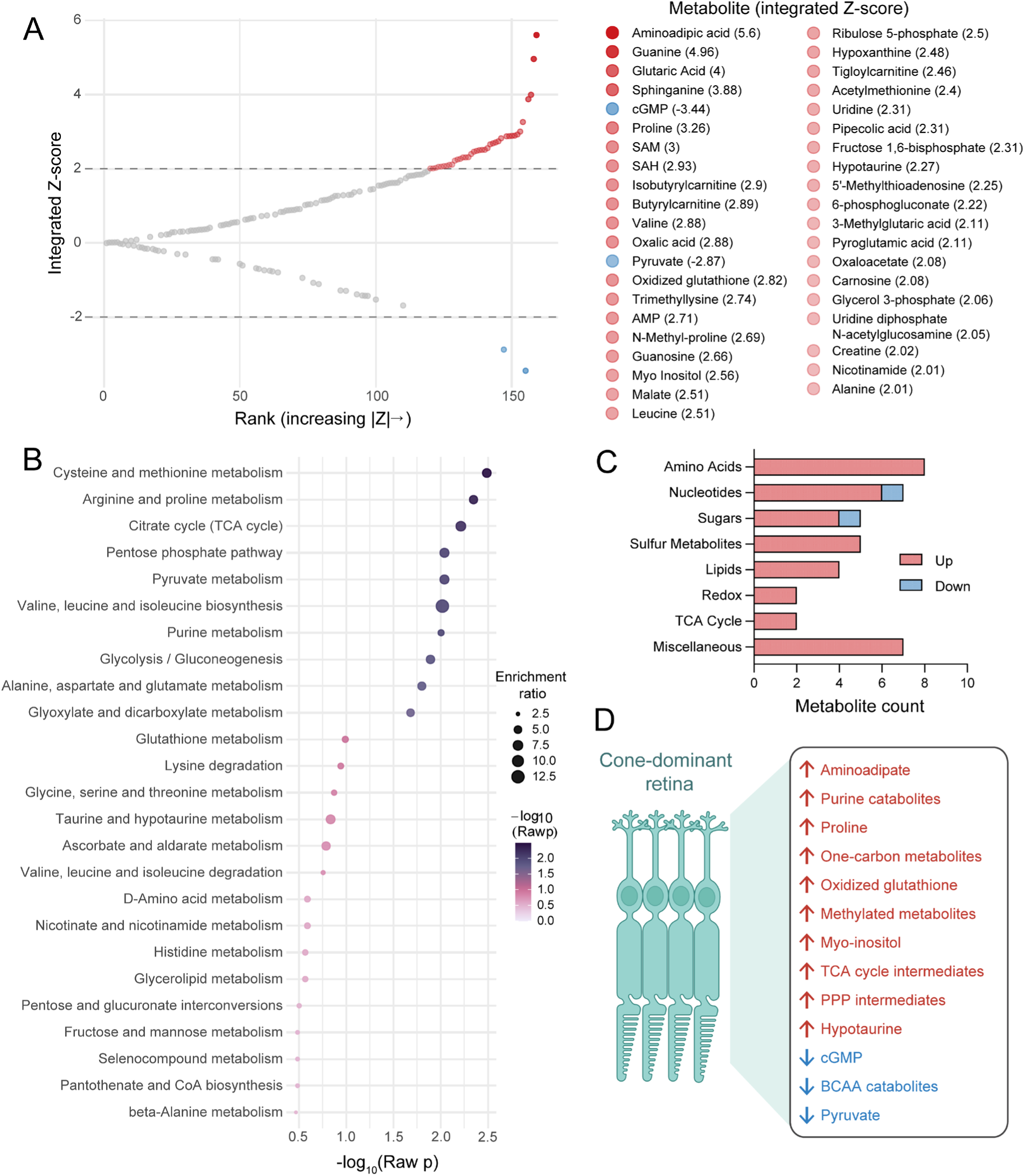
Integrated analysis of retinal metabolite changes associated with relative cone abundance. (A) Scatterplot of metabolites ranked by increasing absolute Z-score, integrated from individual comparison Z-scores using Stouffer’s method. Metabolites exceeding a Z-score threshold of > |2| (equivalent to P < 0.05) are colored red or blue for positive or negative values, respectively. Metabolites exceeding the threshold are listed in order of absolute integrated Z-score magnitude. (B) Metabolic pathway enrichment analysis of integrated retinal metabolite changes with relative cone abundance. Metabolites with an integrated Z-score of > |2| were used for pathway enrichment analysis in MetaboAnalyst 6.0. (C) Metabolite categorization of metabolites with an integrated Z-score of > |2|. (D) Summary diagram of metabolite changes in relatively cone-dominant retinas based on integrated Z-scores of > |2|.

### 2.5. Consistent amino acid and antioxidant changes define cone-dominant RPE metabolism

Similar to our retinal analysis, we integrated Z-scores for each metabolite across the three comparisons performed in RPE/choroid tissues (Table S15). Across models, proline, aminoadipate, and N-methyl-L-proline showed the most consistent increases with higher cone levels. Xanthosine, hypotaurine, trigonelline, and betaine, were consistently elevated, while uracil was significantly decreased (Figure 7A).

**Figure 7.**
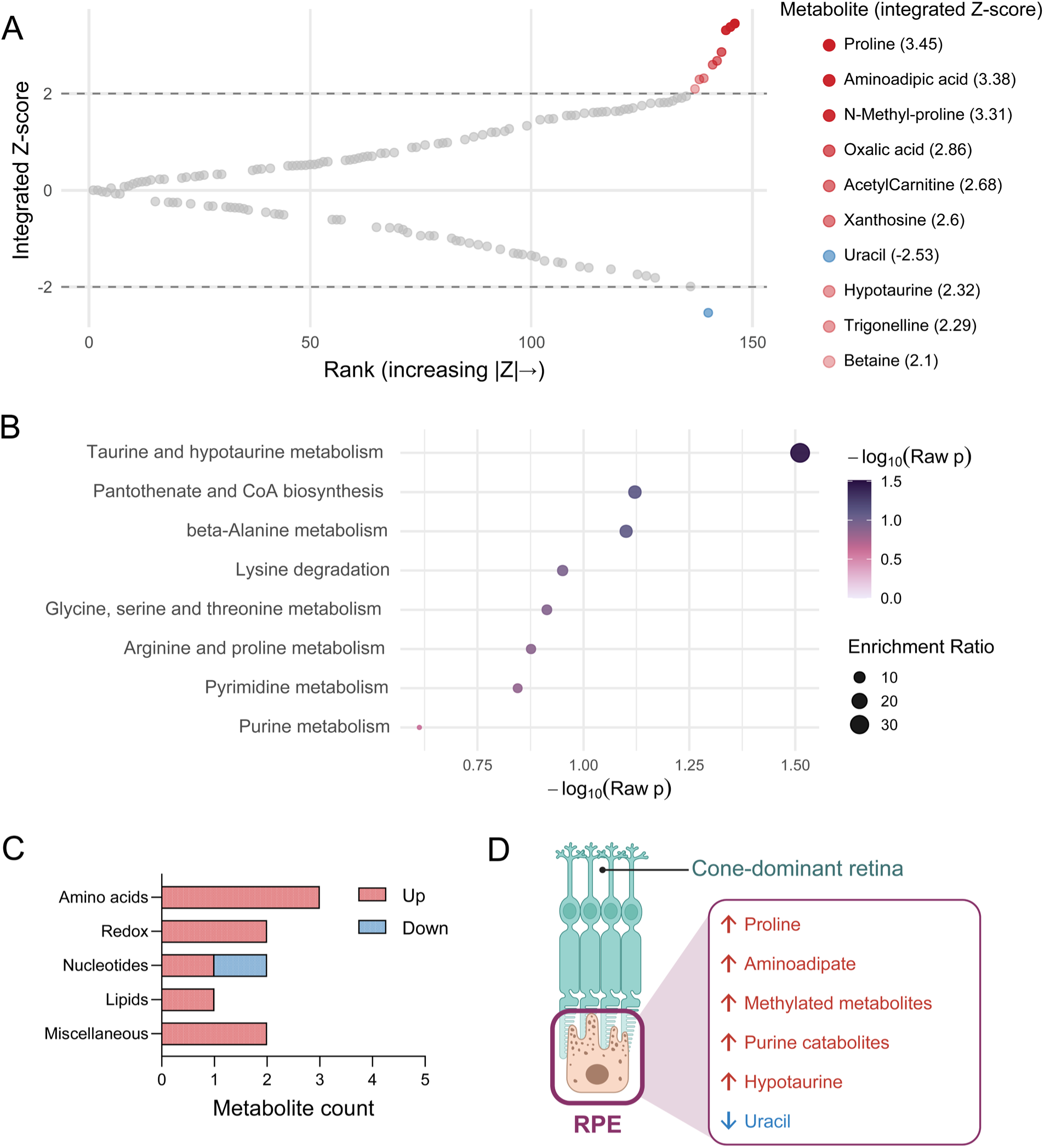
Integrated analysis of RPE/choroid metabolite changes associated with relative cone abundance. (A) Scatterplot of metabolites ranked by increasing absolute Z-score, integrated from individual comparison Z-scores using Stouffer’s method. Metabolites exceeding a Z-score threshold of > |2| (equivalent to P < 0.05) are colored red or blue for positive or negative values, respectively. Metabolites exceeding the threshold are listed in order of absolute integrated Z-score magnitude. (B) Metabolic pathway enrichment analysis of integrated retinal metabolite changes with relative cone abundance. Metabolites with an integrated Z-score of > |2| were used for pathway enrichment analysis in MetaboAnalyst 6.0. (C) Metabolite categorization of metabolites with an integrated Z-score of > |2|. (D) Summary diagram of metabolite changes in RPE/choroids in cone-dominant mice based on integrated Z-scores of > |2|.

Pathway analysis of significantly altered metabolites identified the taurine/hypotaurine pathway as the most strongly affected (Figure 7B), along with pathways involved in alanine, lysine, arginine, and proline metabolism. The amino acid, redox/antioxidant, and nucleotide/nucleoside categories accounted for most of the significantly changed metabolites (Figure 7C). Overall, higher cone proportions were associated with increased hypotaurine, purine catabolism, and one-carbon metabolism in the RPE/choroid, along with altered proline and lysine catabolism (Figure 7D).

## 3. Discussion

Our analysis of cone-altered retina and RPE/choroid metabolomes identified both model-specific and shared metabolic alterations associated with increased cone abundance. Strikingly, cone-dominant retinas showed elevated levels of several amino acids, including proline, valine, leucine, alanine, taurine, hypotaurine, and aminoadipate, as well as higher purine and one-carbon metabolites, alongside lower pyruvate levels. Increased cone abundance was also associated with metabolic changes in the RPE such as higher aminoadipate, proline, and hypotaurine, while proteomic analysis of *Nrl^⁻/⁻^* mice revealed downregulation of multiple key metabolite transporters in the RPE/choroid.

### 3.1. Unique amino acid metabolism in cones

#### 3.1.1. Aminoadipate

The most consistently elevated metabolite associated with cone-dominant retinas was aminoadipate, a central intermediate in lysine metabolism^55^. In humans, lysine is catabolized primarily via the mitochondrial saccharopine pathway or the secondary cytoplasmic and peroxisomal pipecolate pathway, both of which converge at the metabolite aminoadipate semialdehyde^55,56^. Although the RPE is rich in peroxisomes^57,58^, single-cell RNA sequencing (scRNA-seq) data suggest only minimal expression of the key pipecolate pathway enzyme pipecolic acid and sarcosine oxidase (PIPOX) in RPE cells^59^. On the other hand, *ALDH7A1*, which oxidizes aminoadipate semialdehyde to aminoadipate, is highly expressed in the RPE^59,60^. ScRNA-seq data also indicates that cones have low expression of enzymes required for the downstream conversion of aminoadipate to acetyl-CoA, including *AADAT* and *DHTKD1*^56,59^, suggesting they may be unable to catabolize aminoadipate. An alternative source of aminoadipate is via oxidation of lysine residues, which can arise through spontaneous reactions with redox-active metal species or via H_2_O_2_-dependent oxidation by myeloperoxidase^61–63^. This mode of aminoadipate generation has been reported in lysine-rich proteins, such as collagen^62,63^, which is also abundant in retinal structural components such as the inner limiting membrane^64^ and the various layers of retinal vasculature^65,66^. As elaborated below, cones may encounter high oxidative stress relative to rods, perhaps resulting in heightened collagen lysine oxidation to aminoadipate.

Outside the retina, aminoadipate regulates systemic glucose homeostasis, serving as a biomarker of metabolic dysfunction and exerting protective effects in models of obesity and diabetes^67–69^. Specifically, aminoadipate promotes glucose transporter and pyruvate carboxylase expression in pancreatic cells^69^, induces expression of the energy metabolism and mitochondrial regulator PGC-1α^68,70^, and upregulates lipolysis in adipocytes^68^. Moreover, aminoadipate itself is a potentially rich energy source; complete aminoadipate oxidation theoretically generates one FADH_2_ and two molecules each of NADH and acetyl-CoA, corresponding to an estimated yield of ∼24.5 ATP^56,71^ (assuming 2.5 ATP per NADH and 1.5 ATP per FADH_2_). Cones may lack the enzymatic machinery necessary for this oxidation, rendering them unable to use aminoadipate as a fuel source. However, because aminoadipate metabolism in the retina is poorly characterized, this hypothesis requires further experimental validation.

#### 3.1.2. Taurine

Increased cone abundance, particularly in *Nrl^⁻/⁻^* mice, was associated with elevated retinal levels of SAM and hypotaurine. These metabolites, derived from the sulfur-containing amino acids methionine and cysteine, are intermediates in the biosynthesis of taurine, the most abundant amino acid in the retina^72^. Although taurine is present across most retinal cell types, it is most concentrated in the outer nuclear layer, which contains photoreceptor nuclei^73,74^. Importantly, cones are more sensitive than rods to taurine depletion^75,76^. While the precise role of taurine in the retina remains incompletely defined, proposed functions include osmoregulation^77,78^, control of phototransduction and the visual cycle^79,80^, and antioxidant defense^81,82^. In the liver, taurine also regulates cholesterol biosynthesis and excretion^83^, and can conjugate with bile acids to form taurocholate^84^ and taurochenodeoxycholate^85^. Bile acids are also produced in the retina where they confer neuroprotective effects^86,87^. While their cone-specific roles are poorly defined, subcutaneous injection of taurochenodeoxycholate protects against cone loss in a mouse model of retinal degeneration^88^.

Cones have a particularly high sensitivity to oxidative stress^89–91^, which may increase their reliance on abundant taurine to maintain redox homeostasis. Consistently elevated PPP intermediates and hypotaurine, which functions as a free-radical scavenger, support the notion that cones maintain high antioxidant pathway activity^92,93^. Recent investigations using mouse explants indicate that taurine and hypotaurine are shuttled between the retina and RPE^95^, suggesting the RPE supports redox homeostasis in the outer retina. In our study, RPE/choroids from *Nrl^⁻/⁻^* mice showed decreased expression of GAT2, a taurine and hypotaurine transporter^94^. Thus, impaired transport by the RPE may limit taurine availability to the neural retina, necessitating increased endogenous synthesis in cones, as suggested by elevated SAH and hypotaurine levels.

#### 3.1.3. Valine, alanine, leucine and proline

BCAAs and small neutral amino acids including valine, leucine and alanine, were consistently elevated in retinas with high relative cone abundance. Photoreceptor outer segment renewal is a major anabolic sink for amino acids, particularly for opsin synthesis. However, cone outer segment turnover is slower and less organized than in rods^96–99^, potentially reducing anabolic demand. Moreover, valine, leucine and alanine are among the most abundant amino acids in human and mouse rhodopsin^100,101^, suggesting that reduced demand for rod outer segment protein synthesis may contribute to their accumulation. These amino acids are also prominent components of interphotoreceptor matrix (IPM) proteins. The IPM is a specialized extracellular matrix between photoreceptors and the RPE that supports retinal adhesion and metabolite trafficking^102,103^. This scaffold consists of hyaluronan-associated proteoglycans and glycoproteins, including IMPG1, IMPG2, and interphotoreceptor retinoid-binding protein (IRBP)^104^, which are synthesized and degraded primarily by photoreceptors and the RPE^105,106^. Notably, cones express lower levels of key IPM components such as IRBP and IMPG1 compared to rods^59,107^. Reduced synthesis and/or increased turnover of these structural proteins could therefore contribute to elevated valine, alanine, and leucine levels, as these residues are among the most abundant in IRBP^108^ and are also prominent in IMPG1 and IMPG2^109,110^.

Proline was likewise elevated in cone-dominant retinas and is a major constituent of IPM proteins^108–110^. Expression of the proline transporter SLC6A20 was decreased in RPE/choroids of *Nrl^⁻/⁻^* mice. In cultured RPE cells, proline is consumed at exceptionally high rates and metabolized to glutamate, glutamine, aspartate, and other intermediates that support the outer retina^23,24,111^, whereas the retina takes up minimal proline directly^23,24^. Conversely, cone-rich retinal organoids release proline^13^, suggesting that cones may possess the capacity for endogenous proline biosynthesis. These changes suggest a cone-specific metabolic and structural program in the outer retina, with consequences for amino acid flux and IPM homeostasis.

### 3.2. Pyruvate metabolism in cones

The higher energy demands^4^ and greater mitochondrial abundance^6^ indicate that cones rely more heavily on mitochondrial OXPHOS than rods. Consistent with this, we observed lower pyruvate levels alongside increased malate and oxaloacetate in retinas with higher cone abundance, suggesting greater TCA cycle flux. The relative resistance of cones to glycolytic inhibition^9,12^ further implies that they preferentially utilize non-glucose carbon sources including lactate, pyruvate and amino acids. One likely substrate is lactate produced by rods, analogous to lactate utilization by RPE cells^54^. Supporting this notion, cones express higher levels of lactate dehydrogenase B (LDHB) than rods^15^, which favors the conversion of lactate to pyruvate^112^. RPE/choroids showed reduced expression of GLUT1 and MCT1, consistent with altered glucose uptake and lactate transport in cone-dominant retinas. The human macula, which contains a substantially higher density of cones than the peripheral retina, take up more pyruvate uptake in *ex vivo* retinal punches^13^. Similarly, *Nrl^⁻/⁻^* retinal explants demonstrate increased pyruvate uptake compared to wild-type retinas^13^. Pyruvate can also be generated from alternative substrates, including serine^113,114^, alanine^115^, tyrosine^115,116^, glycerol^117^, and TCA intermediates such as malate and oxaloacetate^115^. The accumulation of BCAAs, aminoadipate, and proline may therefore reflect limited capacity of cones to fully oxidize these substrates to acetyl-CoA^118^, increasing reliance on exogenous pyruvate to sustain TCA cycle activity.

### 3.3. Limitations and future directions

As with all genetic and chemically induced models, the methods used to alter cone levels in this study carry model-specific off-target effects. For example, although *Nrl^−/−^* photoreceptors possess many cone-like molecular, morphological, and functional features^44^, *Nrl^-/-^*retinas develop structural abnormalities such as Rosette formation^119–121^, and show increased susceptibility to damage^122^. Similarly, while expression of Cnga3 is cone-specific, *Cnga3^-/-^* mice also undergo progressive secondary rod degeneration and dysfunction^123^. Cnga3 deficiency also leads to metabolic changes directly linked to gene function, such as marked accumulation of cGMP^124^, which was also observed in our analysis. Although high doses of T3 are effective at inducing photoreceptor apoptosis, thyroid hormone signaling influences central metabolic pathways, including those regulating mitochondrial function and lipid metabolism^125,126^. Indeed, T3 profoundly alters the retinal transcriptome, impairing mitochondrial bioenergetics and phototransduction pathways^53^. While a major strength of our approach lies in integrating metabolic changes from multiple distinct models, each with higher cone proportions as the common variable, large model-specific effects (e.g. cGMP accumulation) may be disproportionately over-represented. As such, interpretation of individual metabolite changes should consider the underlying context of each model.

Future studies should continue to characterize the metabolic distinctions between rods and cones by integrating metabolomic, transcriptomic, and proteomic analyses across models of altered cone density. Although our findings indicate differences in lipid biosynthesis and oxidation between rods and cones, a comprehensive lipidomic characterization is still needed, particularly given the central role of disrupted lipid metabolism in AMD and numerous IRDs^127,128^. Our work also highlights BCAAs, proline, taurine, and aminoadipate metabolic pathways as especially prominent in cone cells. Further investigations are warranted to determine how these pathways contribute to cone resilience or vulnerability, RPE-cone metabolic co-operation, and whether targeting them can protect cones from degeneration and functional decline in disease settings. As the field continues to refine cell-specific metabolic mapping, integrating multi-omics data with functional and *in vivo* validation will be essential for translating these insights into effective interventions.

## 4. Methods

### 4.1. Animals

All animal experiments were conducted in accordance with National Institute of Health guidelines, animal protocols approved by the West Virginia University Institutional Animal Care and Use Committee (IACUC), and animal protocols approved by the University of Oklahoma Health Campus IACUC. *Nrl^-/-^*mouse line in a C57BL/6 background^41^ was provided by Dr. Anand Swaroop (National Eye Institute, Bethesda, MD). *Cnga3^-/-^* mouse line in a C57BL/6 background was provided by Dr. Martin Biel (Ludwig-Maximilians-University Munich, Munich)^49,129^. *Nrl^-/-^ Cnga3^-/-^* double knockout mice were generated by cross-mating as previously described^130^.

### 4.2. T3 treatment

All reagents used in this study are listed in Table S16. Mice were administered with T3 via drinking water as described^52^. In short, 10 mg T3 (catalog #T2877, Sigma-Aldrich) was dissolved in 1.0 ml of 1.0N NaOH and diluted in tap water for a final working concentration of 20 µg/mL. *Nrl^-/-^* mother mice were treated with T3 for two weeks, beginning at postnatal day 5 (P5) of the pups. Retinal tissues from the pups were then collected for metabolomic analysis.

### 4.3. Tissue preparation and metabolite extraction

Mouse age and sex were kept consistent within each metabolic comparison. The following genotypes and ages were used: (1) male P30 *Nrl^-/-^* (n=3) and male P30 *Nrl^+/-^* (n=3), (2) male P90 *Cnga3^-/-^* (n=3) and male P90 C57BL/6 (n=3), (3) male P45 *Nrl^-/-^/Cnga3^-/-^* (n=3) and male P45 *Nrl^-/-^* (n=3), (4) female P21 T3-treated *Nrl^-/-^* (n=3) and female P21 *Nrl^-/-^* (n=3). Only retinal tissues were analyzed in the T3-treated comparison. Retinas and RPE/choroid tissues were dissected and snap frozen in liquid nitrogen. Left and right eyes were processed separately, yielding two retina samples and two RPE/choroid samples per mouse. Retinal metabolites were extracted using 80% methanol and homogenized with a microtube homogenizer on dry ice. RPE/choroid tissues were similarly extracted with 80% methanol and homogenized using the Next Advance Bullet BlenderⓇ Gold. Following homogenization, samples were quenched on dry ice for 30 minutes, centrifuged, and supernatants were collected and dried in a vacuum centrifuge (SpeedVac).

### 4.4. LC-MS targeted metabolomic analysis

Dried samples were reconstituted in 40:60 water:acetonitrile. LC-MS/MS was performed using a Shimadzu LC Nexera X2 UHPLC system coupled to a Sciex triple quadrupole QTRAP 5500 mass spectrometer. Chromatographic separation was performed using a 50 mm Waters Acquity UPLC BEH Amide column, with the aqueous mobile phase (A) consisting of water and 10 mM ammonium acetate (pH 8.9), and the organic phase (B) consisting of 95:5 acetonitrile:water with 10 mM ammonium acetate (pH 8.2). All solvents were Fisher Optima LC-MS grade. Metabolites were identified based on accurate mass and retention time. Each sample was analyzed over an 11-minute run using a curved gradient ending in a high-organic mobile phase. Quantification was performed using Sciex MultiQuant 3.0.3 software.

### 4.5. Protein extraction

Retinal and RPE/choroid proteins were extracted using radioimmunoprecipitation assay (RIPA) buffer supplemented with protease inhibitors (10 mL:1 tablet). Samples were homogenized on ice for 15 seconds until the lysate appeared uniformly opaque. Homogenates were rocked on ice for 35 minutes and then centrifuged at 14,000 rpm for 10 min at 4 °C. Supernatants were then collected and frozen at 80℃ until further analysis.

### 4.6. Proteomic analysis

Quantitative proteomic analysis was performed by IDeA National Resource for Quantitative Proteomics (Little Rock, Arkansas). Total protein from each sample was reduced, alkylated, and purified by chloroform/methanol extraction prior to digestion with sequencing grade modified porcine trypsin (Promega). Tryptic peptides were trapped and eluted on 3.5 µm CSH C18 resin (Waters) (4 mm x 75 µm) then separated by reverse phase XSelect CSH C18 2.5 µm resin (Waters) on an in-line 150 x 0.075 mm column using an UltiMate 3000 RSLCnano system (Thermo). Peptides were eluted at a flow rate of 0.300 µL/min using a 60 min gradient from 98% Buffer A (0.1% formic acid, 0.5% acetonitrile):2% Buffer B (0.1% formic acid, 0.5% acetonitrile) to 95:5 at 2.0 minutes to 80:20 at 39.0 minutes to 60:40 at 48.0 minutes to 10:90 at 49.0 minutes and hold until 53.0 minutes and then equilibrated back to 98:2 at 53.1 minutes until 60 minutes. Eluted peptides were ionized by electrospray (2.4 kV) through a heated capillary (275°C) followed by data collection on an Orbitrap Exploris 480 mass spectrometer (Thermo Scientific). Precursor spectra were acquired with a scan from 385-1015 Th at a resolution set to 60,000 with 100% AGC, max time of 50 msec, and an RF parameter at 40%. DIA was configured on the Orbitrap 480 to acquire 50 x 12 Th isolation windows at 15,000 resolution, normalized AGC target 500%, maximum injection time 40 ms). A second DIA was acquired in a staggered window (12 Th) pattern with optimized window placements. Following data acquisition, data were searched using Spectronaut (Biognosys version 19.1) against the UniProt *Mus musculus* database (April 2024) using the directDIA method with an identification precursor and protein q-value cutoff of 1%, generate decoys set to true, the protein inference workflow set to maxLFQ, inference algorithm set to IDPicker, quantity level set to MS2, cross-run normalization set to false, and the protein grouping quantification set to median peptide and precursor quantity. Protein MS2 intensity values were assessed for quality using ProteiNorm^131^. The data was normalized using VSN^132^ and analyzed using proteoDA to perform statistical analysis using Linear Models for Microarray Data (limma) with empirical Bayes (eBayes) smoothing to the standard errors^133,134^. Proteins with an FDR adjusted p-value < 0.05 and a fold change > 2 were considered significant.

### 4.7. Statistical and pathway analysis

Multivariate analysis was performed using a supervised classification model, partial least-squares discriminant analysis (PLS-DA), following Pareto scaling in MetaboAnalyst 6.0 (www.metaboanalyst.ca). In volcano plots, the significance threshold for metabolite comparisons was set at 1.301, corresponding to -log_10_(P < 0.05), as determined using a Student’s unpaired two-tailed T-test. For metabolite classification bar charts, metabolites were manually assigned to categories relevant to retina/RPE physiology. Proteins with significantly altered expression (P < 0.05) in the retina and RPE/choroid between *Nrl^-/-^* and *Nrl^+/-^* mice were curated based on molecular function and biological process gene ontology terms to generate lists of metabolic proteins (Table S11-S12). Integrated pathway analysis of statistically significant proteomic and metabolite changes in this comparison was performed using the joint pathway analysis module on MetaboAnalyst 6.0. Consistency scores for retinal metabolites across comparisons were calculated as a product of support and concordance. Support was defined as the number of comparisons in which the metabolite was significantly altered (Student’s unpaired two-tailed t-test). Concordance was a weighted function capturing the consistency and magnitude of change, defined as:

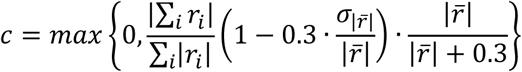

where *r_i_* is the log_2_ fold change in comparison *i*, ∣ *r̄* ∣ is the mean of the absolute fold changes, and *σ*_∣*r*∣_ is the population standard deviation of the absolute fold changes. The coefficient of variation term was set to zero when ∣ *r̄* ∣= 0. Metabolite Z-scores were calculated separately for each comparison and integrated using Stouffer’s method (Table S14-S15). Metabolites with an integrated Z-score > |2| in either retina or RPE/choroid were used for pathway and enrichment analysis in MetaboAnalyst 6.0. Bar charts were generated using GraphPad Prism 10.3.1 (GraphPad Software, Inc). Heatmaps, volcano plots, ranked scatter plots, and metabolite consistency plots were created in R v4.1.2. using the ggplot2 package. The UpSet plot was generated in R using the UpSetR package. Pathway analysis biplots and enrichment plots were visualized ggplot2 using output from MetaboAnalyst 6.0. All explanatory diagrams were created using Biorender (www.biorender.com).

