## Supplemental Figs and Tables S2-S8 and S16 for "Metabolic signatures of cone-dominant and cone-degenerating retinas"

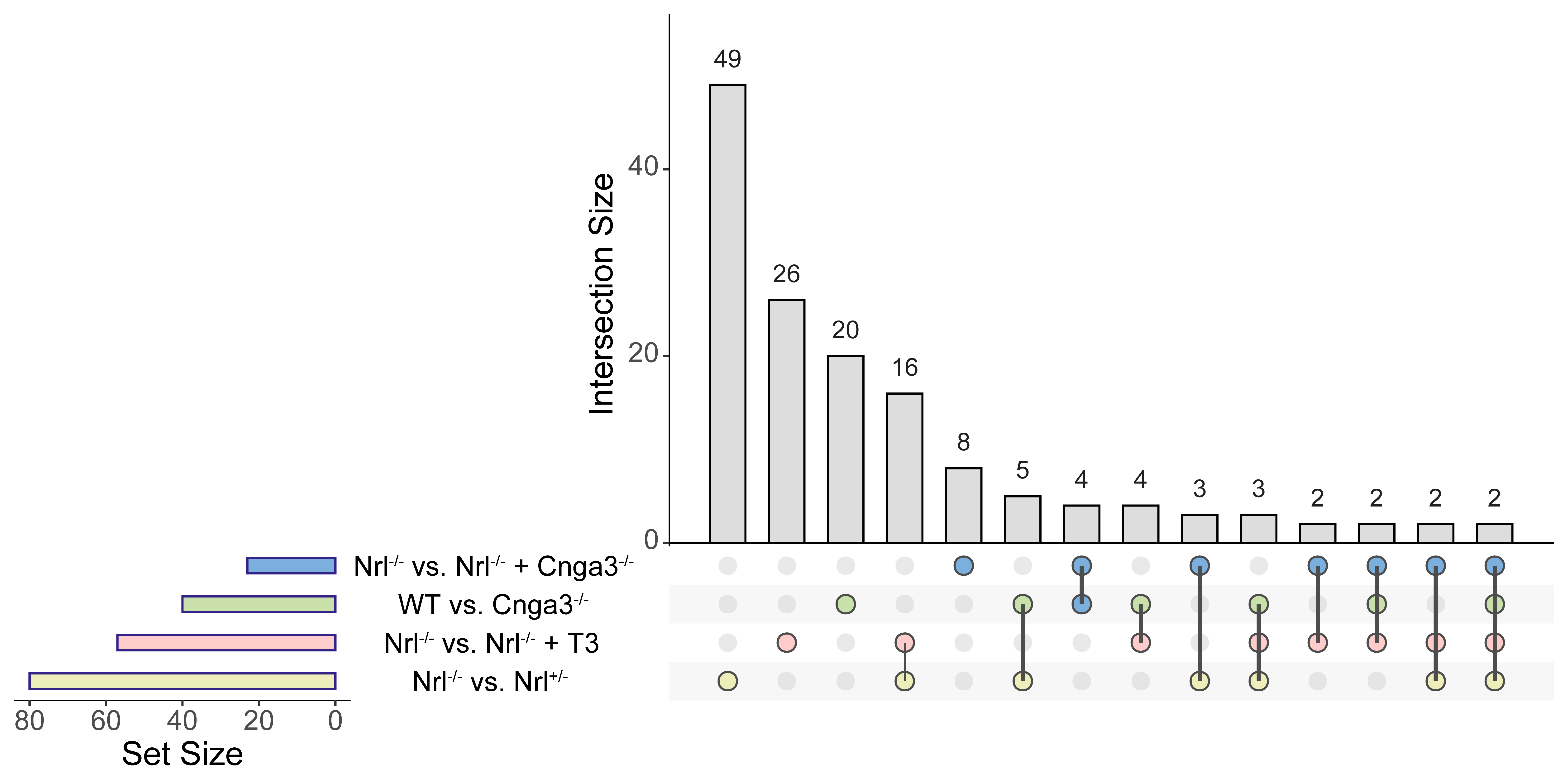
**Figure S1. Shared significant retinal metabolite changes between comparisons.** UpSet plot showing the number of shared metabolites that were significantly changed in each comparison between cone-enriched and cone-depleted retinas. Intersection size refers to the number of significantly changed metabolites in a single comparison or shared between two or more comparisons. Set size is the sum of significantly changed metabolites in a comparison and the number of changed metabolites shared with each of the other comparisons.

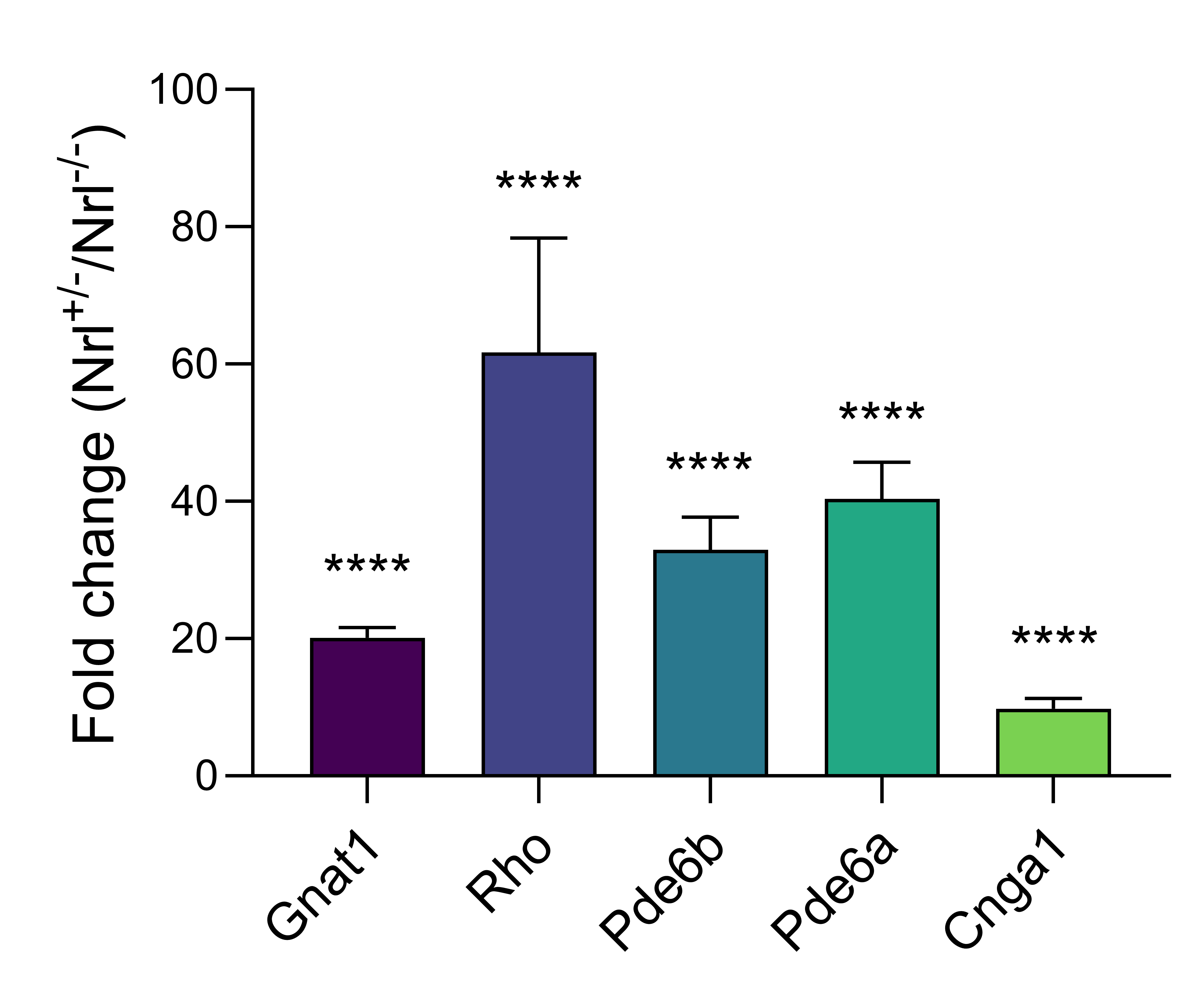

**Figure S2. Relative rod marker protein enrichment in Nrl^+/-^ mice versus Nrl^-/-^ retinas.** Relative protein expression of rod-specific markers in relatively cone-depleted Nrl^+/-^ mice compared to cone-enriched Nrl^-/-^ retinas. Error bars show mean ± SEM. Statistical significance: ****P < 0.0001, determined with unpaired Student’s two-tailed t-test. Gnat1, transducin alpha; Rho, rhodopsin; Pde6b, phosphodiesterase 6B; Pde6a, phosphodiesterase 6A; Cnga1, cyclic nucleotide-gated channel subunit alpha 1.

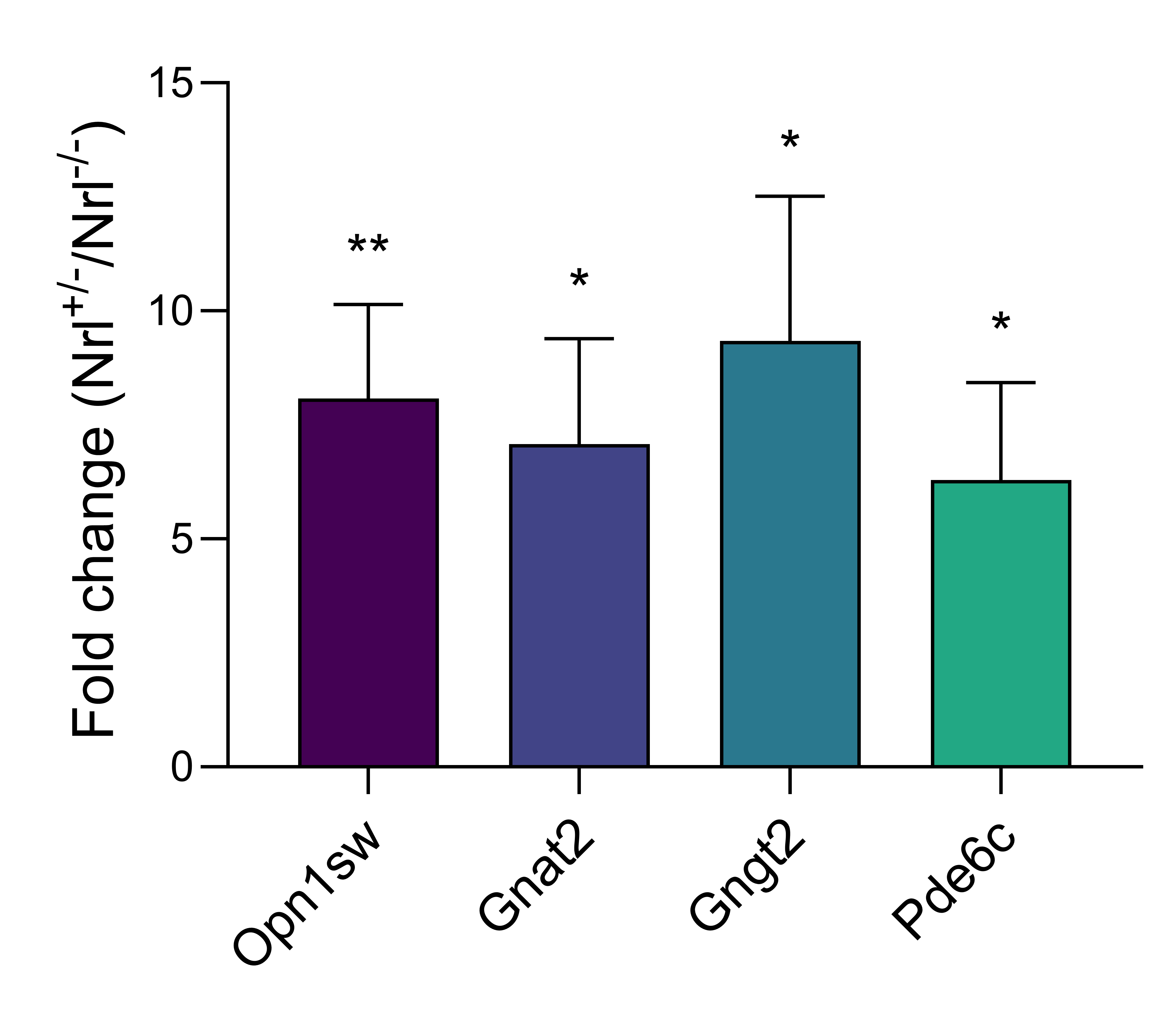

**Figure S3. Relative cone marker protein enrichment in Nrl^-/-^ mice versus Nrl^+/-^ RPE/choroids.** Relative protein expression of cone-specific markers in relatively cone-enriched Nrl^-/-^ mice compared to cone-depleted Nrl^+/-^ RPE/choroids. Error bars show mean ± SEM. Statistical significance: *P < 0.05, **P < 0.01, determined with unpaired Student’s two-tailed t-test. Opn1sw, opsin 1 short wave sensitive; Gnat2, transducin alpha 2; Gngt2, transducin gamma 2; Pde6c, phosphodiesterase 6C.

Table S2. Significantly changed metabolites in Nrl^-/-^ vs. Nrl^+/-^ retinas.

| Metabolite | FC | log2FC | P-value | -log10(P-value) |
| --- | --- | --- | --- | --- |
| Guanine | 0.34 | -1.55 | 0.00003 | 4.471 |
| Aminoadipic acid | 0.38 | -1.41 | 0.00010 | 4.002 |
| Trigonelline | 0.33 | -1.58 | 0.00022 | 3.665 |
| cGMP | 2.04 | 1.03 | 0.00029 | 3.544 |
| Myo Inositol | 0.63 | -0.66 | 0.00083 | 3.080 |
| Acetylmethionine | 0.58 | -0.78 | 0.00131 | 2.883 |
| Threonine | 0.65 | -0.63 | 0.00182 | 2.740 |
| SAH | 0.59 | -0.77 | 0.00423 | 2.374 |
| Nicotinamide | 0.83 | -0.27 | 0.00446 | 2.350 |
| Xanthine | 0.52 | -0.96 | 0.00457 | 2.340 |
| Nicotinamide Riboside | 1.47 | 0.55 | 0.00539 | 2.268 |
| Trimethyllysine | 0.65 | -0.63 | 0.00611 | 2.214 |
| Methylproline | 0.56 | -0.84 | 0.00721 | 2.142 |
| Valine | 0.60 | -0.73 | 0.00859 | 2.066 |
| Pantothenic acid | 0.53 | -0.93 | 0.00887 | 2.052 |
| Hypotaurine | 0.61 | -0.72 | 0.00928 | 2.032 |
| Glutaric Acid | 0.70 | -0.52 | 0.00949 | 2.023 |
| Taurine | 1.39 | 0.47 | 0.00983 | 2.007 |
| Serine | 0.70 | -0.52 | 0.01081 | 1.966 |
| Carnitine | 0.65 | -0.63 | 0.01122 | 1.950 |
| Homoserine | 0.66 | -0.59 | 0.01247 | 1.904 |
| 4-Hydroxyproline | 0.79 | -0.34 | 0.01263 | 1.899 |
| 5'-Methylthioadenosine | 0.65 | -0.62 | 0.01448 | 1.839 |
| Phosphocreatine | 0.62 | -0.68 | 0.01468 | 1.833 |
| Alanine | 0.64 | -0.65 | 0.01502 | 1.823 |
| Malate | 0.81 | -0.30 | 0.01502 | 1.823 |
| 2-Methylbutyroylcarnitine | 0.59 | -0.75 | 0.01529 | 1.816 |
| SAM | 0.61 | -0.70 | 0.01783 | 1.749 |
| Guanosine | 0.75 | -0.42 | 0.01836 | 1.736 |
| Arginine | 0.71 | -0.48 | 0.02060 | 1.686 |
| Pipecolic acid | 0.55 | -0.86 | 0.02096 | 1.679 |
| Fructose 1,6-bisphosphate | 0.71 | -0.50 | 0.02100 | 1.678 |
| Sphinganine | 0.79 | -0.34 | 0.02298 | 1.639 |
| N-alpha-Acetyl-L-lysine | 0.70 | -0.51 | 0.02395 | 1.621 |
| Oxidized glutathione | 0.51 | -0.96 | 0.02614 | 1.583 |
| 6-phosphogluconate | 0.80 | -0.33 | 0.02638 | 1.579 |
| Glutathione | 0.39 | -1.34 | 0.03144 | 1.503 |
| Glutamax | 0.80 | -0.32 | 0.03211 | 1.493 |
| Leucine | 0.78 | -0.36 | 0.03279 | 1.484 |
| Ophthalmic Acid | 0.59 | -0.75 | 0.03333 | 1.477 |
| 3-Methylglutaric acid | 0.73 | -0.46 | 0.03449 | 1.462 |
| Pyroglutamic acid | 0.59 | -0.77 | 0.03508 | 1.455 |
| Oxaloacetate | 0.76 | -0.40 | 0.03734 | 1.428 |
| Adenine | 1.35 | 0.43 | 0.03929 | 1.406 |
| Glycerol 3-phosphate | 0.69 | -0.53 | 0.03945 | 1.404 |
| Proline | 0.68 | -0.55 | 0.04335 | 1.363 |
| Ribulose 5-phosphate | 0.71 | -0.50 | 0.04481 | 1.349 |
| Citrulline | 0.74 | -0.44 | 0.04784 | 1.320 |
| Butyrylcarnitine | 0.67 | -0.58 | 0.04819 | 1.317 |

Table S3. Significantly changed metabolites in Cnga3^-/-^ vs. WT retinas.

| Metabolite | FC | log2FC | P-value | -log10(P-value) |
| --- | --- | --- | --- | --- |
| Butyrylcarnitine | 0.83 | -0.26 | 0.002 | 2.654 |
| Glucose | 1.54 | 0.62 | 0.004 | 2.399 |
| a-ketoglutarate | 1.65 | 0.73 | 0.008 | 2.089 |
| cGMP | 1.52 | 0.61 | 0.010 | 2.017 |
| Cytidine | 0.62 | -0.68 | 0.010 | 2.004 |
| Isobutyrylcarnitine | 0.82 | -0.28 | 0.010 | 1.997 |
| Arginine | 1.22 | 0.29 | 0.011 | 1.945 |
| Aminoadipic acid | 0.44 | -1.17 | 0.012 | 1.927 |
| UDP-Glucose | 1.48 | 0.57 | 0.015 | 1.811 |
| cAMP | 1.64 | 0.72 | 0.025 | 1.601 |
| Propionylcarnitine | 0.68 | -0.55 | 0.030 | 1.523 |
| Phosphocreatine | 2.64 | 1.40 | 0.031 | 1.514 |
| Betaine | 0.81 | -0.30 | 0.036 | 1.448 |
| Tigloylcarnitine | 0.86 | -0.21 | 0.038 | 1.424 |
| Ornithine | 0.72 | -0.47 | 0.040 | 1.393 |
| ATP | 2.92 | 1.55 | 0.041 | 1.385 |
| AcetylCarnitine | 0.78 | -0.36 | 0.042 | 1.381 |
| FAD | 1.59 | 0.67 | 0.043 | 1.368 |
| Uridine | 0.53 | -0.91 | 0.045 | 1.347 |
| ADP | 1.45 | 0.54 | 0.047 | 1.327 |

Table S4. Significantly changed metabolites in Nrl^-/-^ vs. Cnga3^-/-^ + Nrl^-/-^ retinas.

| Metabolite | FC | Log2 FC | P-value | -log10 P-value |
| --- | --- | --- | --- | --- |
| cGMP | 149.33 | 7.22 | 0.005 | 2.31 |
| Fructose | 4.68 | 2.23 | 0.008 | 2.11 |
| 1-Methyladenosine | 1.92 | 0.94 | 0.010 | 2.02 |
| N1-Methylnicotinamide | 1.23 | 0.30 | 0.012 | 1.92 |
| Riboflavin | 2.19 | 1.13 | 0.032 | 1.49 |
| Glutamax | 1.84 | 0.88 | 0.038 | 1.42 |
| Tyrosine | 1.40 | 0.48 | 0.042 | 1.37 |
| Aminoadipic acid | 0.40 | -1.33 | 0.047 | 1.33 |

Table S5. Significantly changed metabolites in Nrl^-/-^ vs. Nrl^-/-^ + T3 retinas.

| Metabolite | FC | Log2 FC | P-value | -log10 P-value |
| --- | --- | --- | --- | --- |
| Malate | 0.78 | -0.36 | 0.0004 | 3.35 |
| Sphinganine | 0.32 | -1.65 | 0.0013 | 2.87 |
| Aminoadipic acid | 0.19 | -2.40 | 0.0049 | 2.31 |
| Arginine | 0.47 | -1.09 | 0.0073 | 2.13 |
| AMP | 0.63 | -0.67 | 0.0076 | 2.12 |
| Uracil | 2.32 | 1.21 | 0.0080 | 2.10 |
| Guanine | 0.42 | -1.25 | 0.0094 | 2.03 |
| Oxidized glutathione | 0.55 | -0.86 | 0.0103 | 1.99 |
| ADP | 0.60 | -0.75 | 0.0113 | 1.95 |
| Oxalic acid | 0.50 | -1.01 | 0.0139 | 1.86 |
| XMP | 0.55 | -0.87 | 0.0142 | 1.85 |
| Argininosuccinic acid | 0.63 | -0.66 | 0.0208 | 1.68 |
| Glutaric Acid | 0.52 | -0.94 | 0.0246 | 1.61 |
| Proline | 0.14 | -2.82 | 0.0254 | 1.60 |
| Ribulose 5-phosphate | 0.75 | -0.41 | 0.0292 | 1.53 |
| GMP | 0.56 | -0.84 | 0.0297 | 1.53 |
| cGMP | 0.54 | -0.88 | 0.0319 | 1.50 |
| Creatine | 0.56 | -0.84 | 0.0330 | 1.48 |
| Leucine | 0.52 | -0.95 | 0.0332 | 1.48 |
| Creatinine | 0.65 | -0.61 | 0.0343 | 1.47 |
| Guanosine | 0.68 | -0.56 | 0.0356 | 1.45 |
| Carnitine | 0.72 | -0.48 | 0.0361 | 1.44 |
| Glucose 6-phosphate | 0.73 | -0.45 | 0.0371 | 1.43 |
| Valine | 0.37 | -1.43 | 0.0402 | 1.40 |
| Acetylmethionine | 0.45 | -1.14 | 0.0420 | 1.38 |
| SAM | 0.46 | -1.12 | 0.0494 | 1.31 |

Table S6. Significantly changed metabolites in Nrl^-/-^ vs. Nrl^+/-^ RPE/choroids.

| Metabolite | FC | log2FC | P-value | -log10 P-value |
| --- | --- | --- | --- | --- |
| Thiamine | 0.61 | -0.72 | 0.0007 | 3.16 |
| Methylproline | 1.94 | 0.96 | 0.0009 | 3.03 |
| Riboflavin | 1.67 | 0.74 | 0.0033 | 2.48 |
| Oxalic acid | 1.39 | 0.47 | 0.0042 | 2.37 |
| 2-Methylbutyroylcarnitine | 1.72 | 0.78 | 0.0069 | 2.16 |
| Hypotaurine | 2.46 | 1.30 | 0.0070 | 2.15 |
| Valine | 1.47 | 0.56 | 0.0090 | 2.05 |
| Xanthosine | 1.63 | 0.70 | 0.0094 | 2.03 |
| Histamine | 1.55 | 0.63 | 0.0142 | 1.85 |
| Creatinine | 1.29 | 0.37 | 0.0163 | 1.79 |
| Myo Inositol | 1.38 | 0.47 | 0.0204 | 1.69 |
| Leucine | 1.39 | 0.48 | 0.0264 | 1.58 |
| Alanine | 1.42 | 0.51 | 0.0265 | 1.58 |
| Isobutyrylcarnitine | 1.50 | 0.59 | 0.0277 | 1.56 |
| Butyrylcarnitine | 1.49 | 0.58 | 0.0281 | 1.55 |
| Aspartic acid | 2.23 | 1.15 | 0.0338 | 1.47 |
| Glutamic acid | 2.62 | 1.39 | 0.0489 | 1.31 |

Table S7. Significantly changed metabolites in Cnga3^-/-^ vs. WT RPE/choroids.

| Metabolite | FC | Log2 FC | P-value | -log10 P-value |
| --- | --- | --- | --- | --- |
| Thiamine | 0.68 | -0.55 | 0.002 | 2.71 |
| AcetylCarnitine | 0.31 | -1.68 | 0.002 | 2.70 |
| IPP | 1.29 | 0.36 | 0.003 | 2.56 |
| Carnosine | 1.79 | 0.84 | 0.003 | 2.46 |
| Riboflavin | 0.71 | -0.49 | 0.004 | 2.40 |
| Citraconic Acid | 1.81 | 0.86 | 0.007 | 2.18 |
| Uracil | 1.49 | 0.57 | 0.010 | 2.02 |
| Succinate | 1.22 | 0.29 | 0.015 | 1.83 |
| Proline | 0.76 | -0.39 | 0.016 | 1.81 |
| Butyrylcarnitine | 1.44 | 0.52 | 0.018 | 1.73 |
| Aminoadipic acid | 0.42 | -1.25 | 0.026 | 1.59 |
| Isobutyrylcarnitine | 1.46 | 0.54 | 0.027 | 1.57 |
| Myristoylcarnitine | 1.24 | 0.31 | 0.029 | 1.54 |
| Acetylalanine | 1.24 | 0.32 | 0.039 | 1.40 |
| Argininosuccinic acid | 1.51 | 0.59 | 0.040 | 1.40 |
| cAMP | 1.97 | 0.98 | 0.041 | 1.39 |
| Aconitate | 1.49 | 0.58 | 0.041 | 1.39 |
| Tryptophan | 0.78 | -0.37 | 0.044 | 1.35 |
| Myo Inositol | 1.20 | 0.26 | 0.045 | 1.35 |

Table S8. Significantly changed metabolites in Nrl^-/-^ vs. Cnga3^-/-^ + Nrl^-/-^ RPE/choroids.

| Metabolite | FC | Log2FC | P-value | -log10 P-value |
| --- | --- | --- | --- | --- |
| Pantothenic acid | 1.47 | 0.55 | 0.003 | 2.55 |
| Riboflavin | 1.14 | 0.19 | 0.007 | 2.13 |
| Guanine | 1.62 | 0.69 | 0.011 | 1.95 |
| cGMP | 2.80 | 1.48 | 0.020 | 1.70 |
| Proline | 0.83 | -0.28 | 0.028 | 1.55 |
| Inosine | 1.73 | 0.79 | 0.044 | 1.36 |
| Kynurenine | 0.81 | -0.31 | 0.044 | 1.36 |
| Heptadecanoic acid | 3.97 | 1.99 | 0.048 | 1.32 |

Table S16. Reagents and materials used in this study.

| Reagent | Catalog | Company | Location |
| --- | --- | --- | --- |
| RIPA Lysis and Extraction Buffer | 89900 | Thermo scientific | Rockford IL USA |
| Pierce Protease and Phosphatase Inhibitor Mini Tablets | A32959 | Thermo scientific | Rockford IL USA |
| Pierce™ BCA Protein Assay Kit | 23225 | Thermo scientific | Rockford IL USA |
| Phosphate buffer saline | 10010 | Thermo scientific | Rockford IL USA |
| Methanol | A456 | Fisher Scientific | Pittsburgh PA USA |
| HPLC Water | W6 | Fisher Scientific | Pittsburgh PA USA |
| Acetonitrile | A955 | Fisher Scientific | Pittsburgh PA USA |
| Ammonium Acetate | 431311 | Sigma Aldrich | Saint Louis MO USA |
| Ammonium Hydroxide | 338818 | Sigma Aldrich | Saint Louis MO USA |
| Nicotinamide (D4, 98%) | DLM-6883 | Cambridge Isotope Laboratories | Tewksbury MA USA |
| Accquity UPLC BEH Amide Column, 130Å, 1.7 µm, 2.1 mm X 150 mm | 186004802 | Waters | Milford MA USA |
